# SpatialSPM: Statistical parametric mapping for the comparison of gene expression pattern images in multiple spatial transcriptomic datasets

**DOI:** 10.1101/2023.06.26.546605

**Authors:** Jungyoon Ohn, Mi-Kyoung Seo, Jeongbin Park, Daeseung Lee, Hongyoon Choi

**Author notes:** **Corresponding author:** Hongyoon Choi, MD., PhD., Department of Nuclear Medicine, Seoul National University Hospital, Seoul, Republic of Korea; Department of Nuclear Medicine, Seoul National University College of Medicine, Republic of Korea. These authors contributed equally to this work (Jungyoon Ohn, Mi-Kyoung Seo).

## Abstract

Spatial transcriptomic (ST) techniques help us understand the gene expression levels in specific parts of tissues and organs, providing insights into their biological functions. Even though ST dataset provides information on the gene expression and its location for each sample, it is challenging to compare spatial gene expression patterns across tissue samples with different shapes and coordinates. Here, we propose a method that reconstructs ST data into multi-dimensional image matrices to ensure comparability across different samples through spatial registration process. We demonstrated the applicability of this method by using two mouse brain ST datasets to investigate and directly compare gene expression in a specific anatomical region of interest, pixel by pixel, across various biological statuses. It can produce statistical parametric maps to find specific regions with differentially expressed genes across tissue samples. Our approach provides an efficient way to analyze ST datasets and may offer detailed insights into various biological conditions.

## Introduction

Spatial transcriptomics (ST) has emerged as a powerful technique for obtaining location-specific transcriptomic information in tissues and organs ^1-3^. This technique provides valuable insights into biological physiology, particularly regarding spatial heterogeneity. Therefore, investigating transcriptomic alterations in ST under pathological conditions, such as injury, inflammatory diseases, or cancer, can enable researchers to uncover the underlying pathogenesis specific to each detailed segment of the organ ^4-6^. However, the ST dataset itself only provides information about the gene expression level and relative location of each transcript, without any actual anatomical information ^1^. Furthermore, due to the different shapes and morphologies of each tissue sample for ST dataset, direct comparisons among them become problematic ^3^. These limitations make it challenging to conduct research solely based on ST dataset.

To overcome the limitations of ST dataset, researchers often incorporate additional information, such as H&E staining images, immunofluorescent staining images based on cell markers, or single-cell RNA sequencing datasets ^1,7^. These complementary sources of information enable a comprehensive understanding of the spatial gene expression patterns within organs. Nevertheless, comparing multiple ST datasets for a specific region of interest (ROI) can be challenging, particularly when dealing with organs that have complex anatomical structures, such as the brain ^8^.

In this regard, we propose a new approach called SpatialSPM **(Fig. 1)**, which reconstructs gene expression pattern images based on ST dataset and incorporates a spatial registration process for the images. This approach allows us to provide gene expression levels in specific anatomical ROIs and generate statistical parametric maps at a pixel level. Furthermore, it enables direct comparison of gene expression in specific anatomical ROIs across various biological statuses, providing valuable insights into underlying pathogenesis.

## Results

### SpatialSPM for the pixel-wise comparative analysis in spatially registered images based on the ST dataset

The overall workflow of SpatialSPM is depicted in **Fig. 1** (please refer to the Materials and Methods section for more details). This method utilized a convolutional operation to estimate gene expression values for locations between the detection spots in ST dataset by superimposing Gaussian values from neighboring spots. The resulting two-dimensional (2D) images for each gene were compiled and loaded as matrices. Next, score maps of highly-variable genes (HVGs) were constructed, which are used to generate a transformation matrix. This transformation matrix was then applied to all genes in the 2D image matrices, resulting in transformed matrices, which is spatial registration process. Comparative analysis could be performed at the pixel level across the spatially registered images to evaluate gene expression levels in specific brain regions.

**Figure 1.**
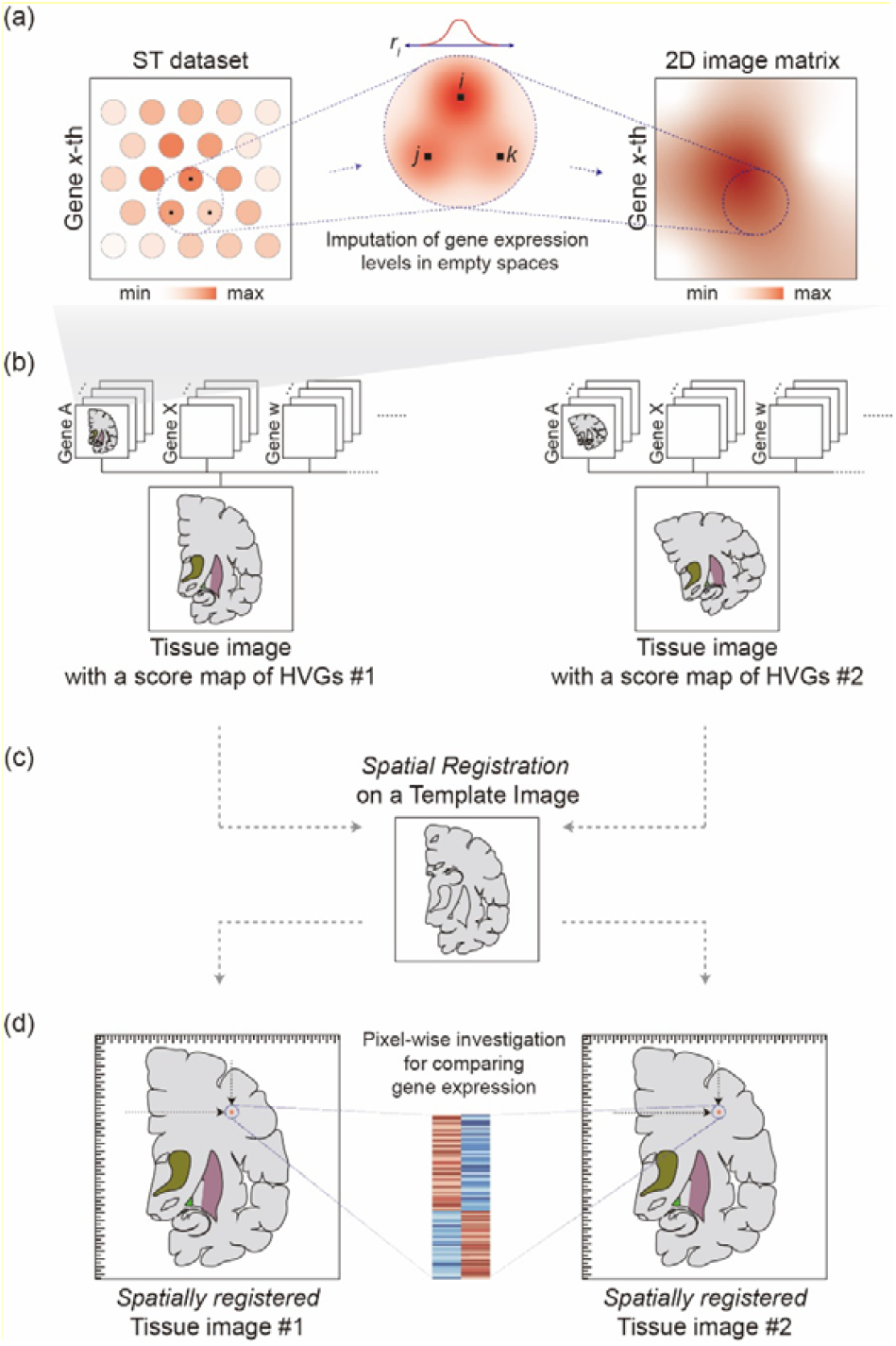
SpatialSPM: Pixel-wise comparative analysis in *‘spatially registered’* images generated based on the ST dataset. (a) Generation of a two-dimensional expression image matrix by imputing the gene expression levels in empty spaces between the detection spots. (b) Identification of gene expression patterns for constructing a tissue image with a score map of HVGs for the spatial registration process (c). (d) Pixel-wise investigation for comparing gene expression levels in a specific anatomical region of the spatially registered images. **Abbreviation**: HVGs, highly variable genes; ST, spatial transcriptomics

### Investigation of the intracranial hemorrhage mouse model brain ST dataset by SpatialSPM

Our SpatialSPM methodology was applied to ST datasets obtained from mouse animal models with intracranial hemorrhage, induced by Heme-Albumin injection into the striatum ^9^. Five ST datasets were processed into multiple 2D image matrices of gene expression values, representing brain tissues from the Sham group and Heme-Albumin injection groups with concentrations of 0.03nM, 1.25nM, 5.0nM, and 10.0nM. HVGs score maps were also constructed for each group (**Fig. 2a**). The HVGs score map exhibited clear distinctions based on the specific brain anatomical structure in each group. Subsequently, the images were processed by the spatial registration process, using the image of the sham group as the template (**Fig. 2b)**. As a result of this process, five spatially registered 2D brain images were obtained, exhibiting identical contour and internal arrangement **(Fig. 2c**).

**Figure 2.**
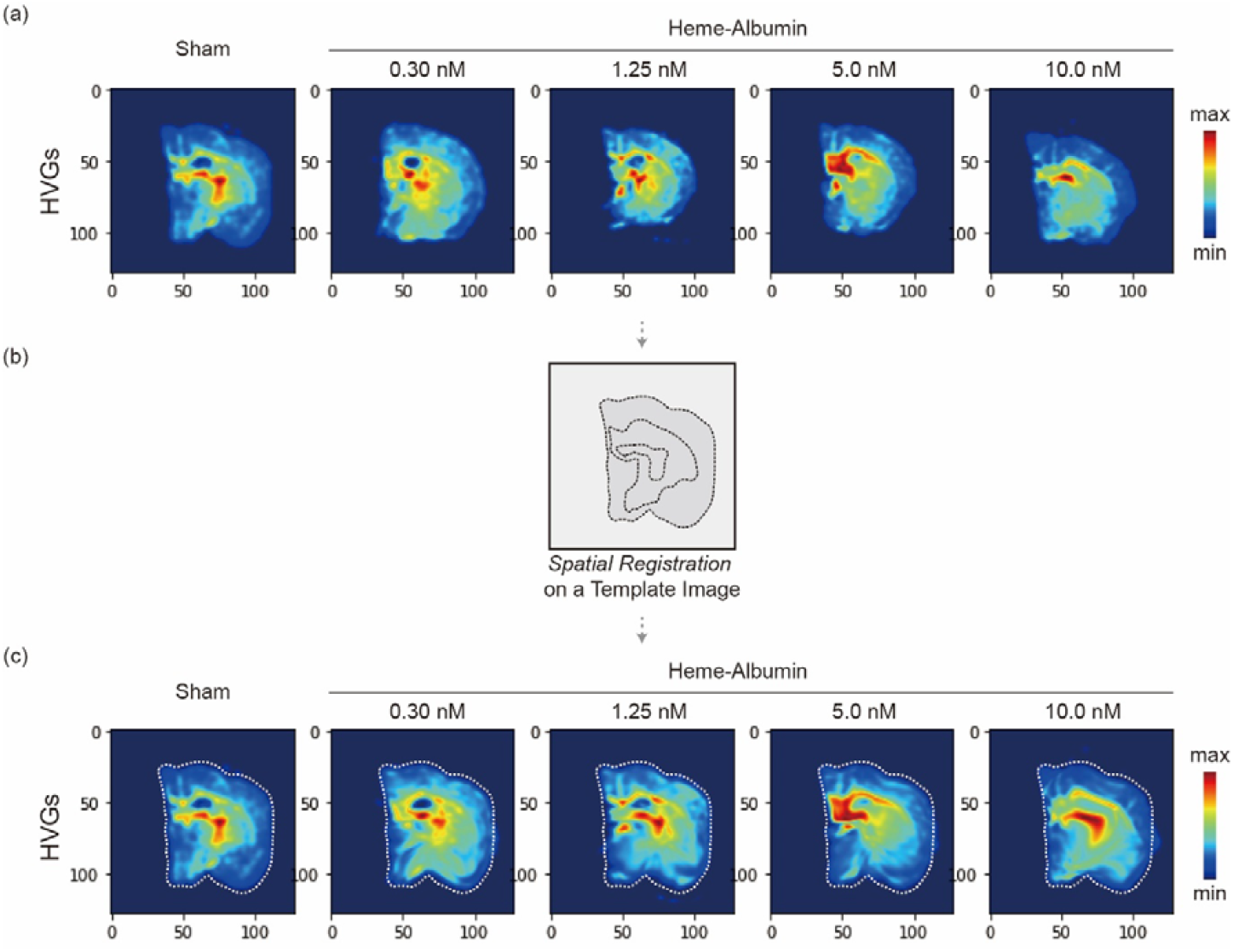
Investigation of the intracranial hemorrhage mouse model brain ST dataset by SpatialSPM. (a) Five ST datasets were obtained from five groups: Sham, 0.03nM, 1.25nM, 5.0nM, and 10.0nM of Heme-Albumin injection groups. Each dataset was processed into a 2D image matrix representing gene expression values. The HVGs score maps was constructed based on these matrices. (b) The spatial registration process resulted in five spatially registered 2D brain images (c). **Abbreviation**: HVGs, highly-variable genes; ST, spatial transcriptomics; 2D, two-dimensional;

In five spatially registered 2D brain images, glial fibrillary acidic protein (GFAP) gene (*Gfap*) expression level was displayed as a demonstrative example of a specific gene (**Fig. 3a**). GFAP is a signature intermediate filament of astrocytes and its expression level is indicative of the severity and extent of intracranial pathologic status after traumatic brain injury ^10^. The expression level of *Gfap* in striatum and cortex of brain was tightly correlated with the Heme-Albumin concentration. Additionally, the pixel-wise correlation analysis was performed and represented as a parametric map showing Pearson’s correlation coefficients for each gene (**Fig. 3b**). Notably, the expression level of *Gfap* in the center of striatum (Heme-Albumin injection site) showed statistically less correlated with the concentration, suggesting that *Gfap* expression levels in the injection site were not affected by Heme-Albumin concentration.

**Figure 3.**
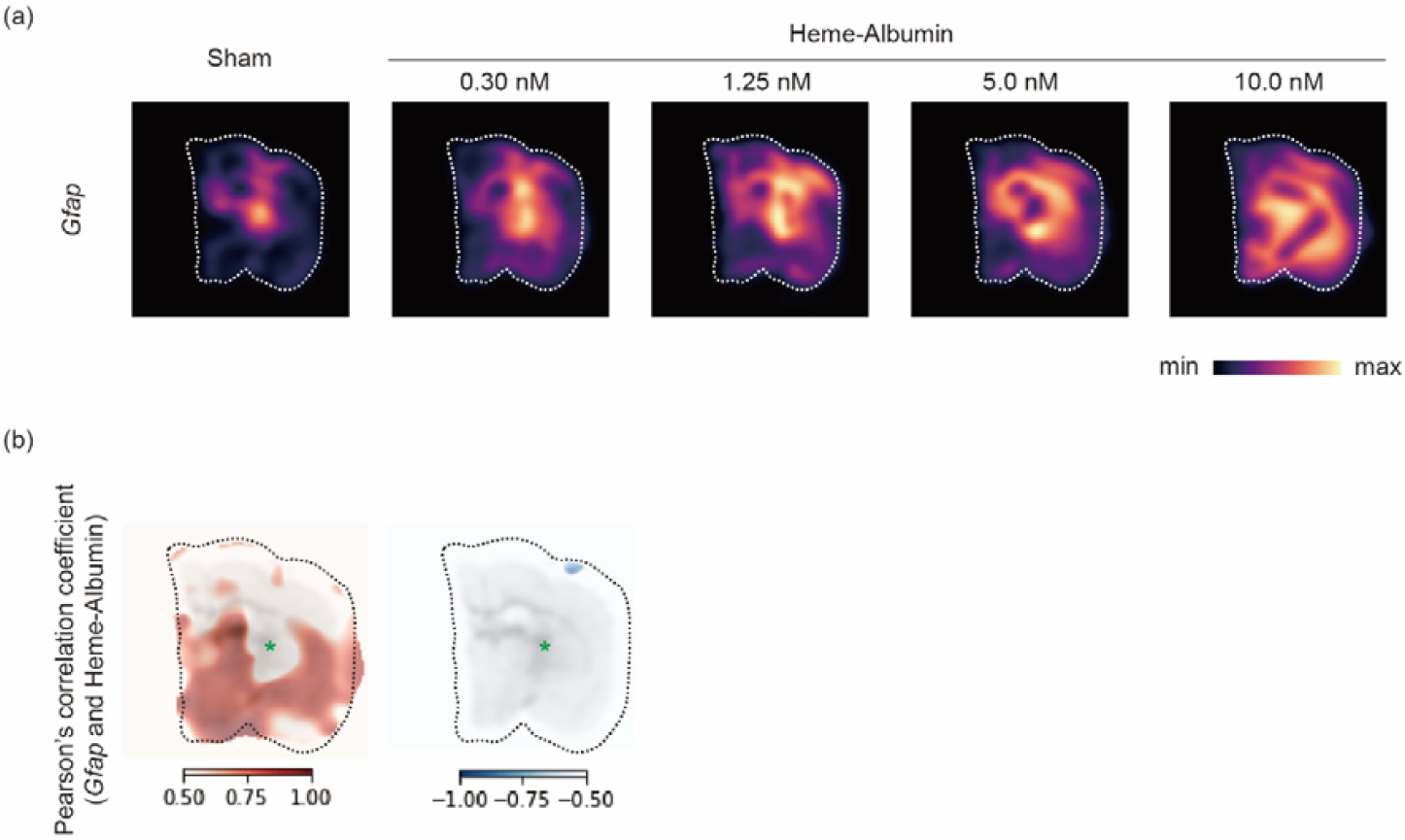
*Gfap* expression levels in spatially registered brain images of intracranial hemorrhage mouse model. (a) Heatmaps illustrated based on *Gfap* expression levels across five groups (Sham, 0.03nM, 1.25nM, 5.0nM, and 10.0nM of Heme-Albumin injection groups). (b) Heatmaps illustrated based on the Pearson’s correlation coefficient between *Gfap* expression and Heme-Albumin concentration (asterisk: Heme-Albumin injection site). **Abbreviation**: *Gfap*, glial fibrillary acidic protein;

Also, parametric maps can be used to display and statistically analyze all other gene features in the ST datasets. **Supplementary Fig. 1** shows the representative parametric maps of the top 30 genes that were highly correlated (Pearson’s correlation coefficient > 0.3) with Heme-Albumin concentration.

### Investigation of PSAPP mouse model of Alzheimer’s disease brain ST dataset by SpatialSPM

SpatialSPM can also be applied for comparative analysis between two groups. Total six ST datasets from the amyloid-depositing Alzheimer’s disease mouse model (*APP*^*KM670/671NL*^*/PSEN1*^Δ*exon9*^; *PSAPP*) with conditional *Inpp5d* knockdown in microglia (*PSAPP*/*Inpp5d*^fl/fl^/*Cx3xr1*^CreER+^) were utilized ^11^. Among them, three datasets were acquired from corn oil (CO) treated mice (PSAPP-CO) and the other datasets were from tamoxifen (TAM) treated mice (PSAPP-TAM) with down-regulation of *Inpp5d* in microglia.

The ST datasets were processed into 2D image matrix of each gene expression value, constructing the HVGs score map images, respectively **(Fig. 4a)**. Despite all the maps being images of coronal brain tissue sections, direct comparison and analysis of gene expression status in specific anatomical locations was challenging due to variations in image morphologies. The spatial registration process of the images on a template image (in this case, the first image of the PSAPP-CO group) was performed (**Fig. 4b**), resulting six spatially registered 2D brain images with same configuration (**Fig. 4c**). The scores of HVGs were clearly distinguishable according to the brain anatomic structure.

**Figure 4.**
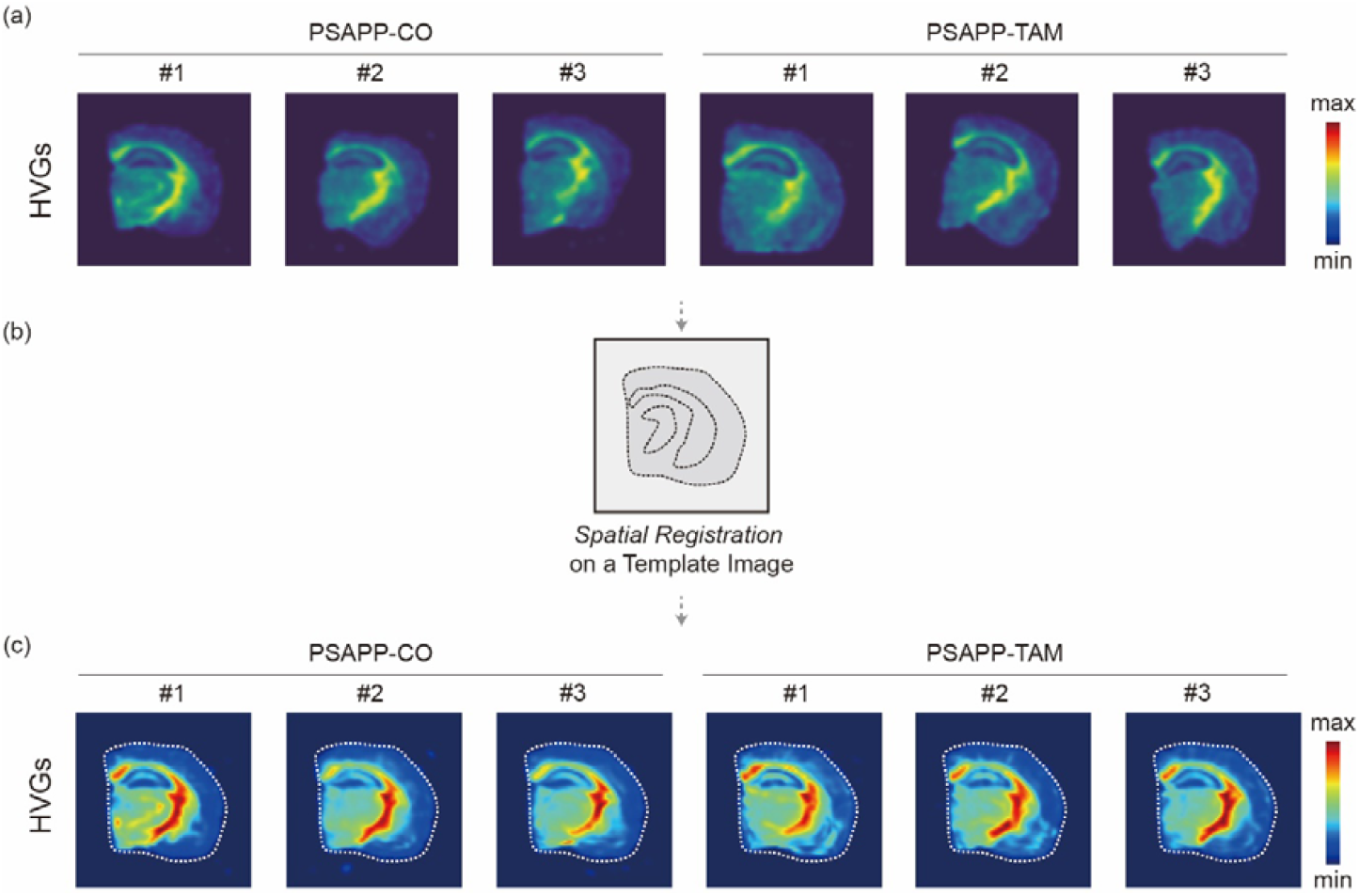
Investigation of the Alzheimer’s disease mouse model brain ST dataset by SpatialSPM. (a) Six ST datasets were obtained from two groups: PSAPP-CO and PSAPP-TAM. Each dataset was processed into 2D image matrix representing each gene expression value. The HVGs score maps were constructed based on these matrices. (b) The spatial registration process resulted in Six spatially registered 2D brain images (c). **Abbreviation**: CO, corn oil; HVGs, highly-variable genes; PSAPP, APP^KM670/671NL^/PSEN1^Δexon9^; ST, spatial transcriptomics; TAM, tamoxifen; 2D, two-dimensional;

After the spatial registration process, *Cst7* gene, a well-known biomarker of neuro-inflammation related to the amyloid beta plaque in Alzheimer’s disease, was selected as an example for investigation and the gene expression levels were shown in the images **(Fig. 5a)**. The expression level of *Cst7* was higher in the brain of the PSAPP-TAM group compared to the PSAPP-CO group. The pixel-wise T-test was performed between two groups (PSAPP-CO and PSAPP-TAM with three biological replicates in each group) for mapping the statistics (presented as *T-*score) for *Cst7* gene expression alteration in each anatomic location according to the conditional *Inpp5d* knockdown in brain microglia **(Fig. 5b)**. It was found that *Cst7* expression was elevated in striatal and cortex area of PSAPP-TAM brain with statistical significance.

**Figure 5.**
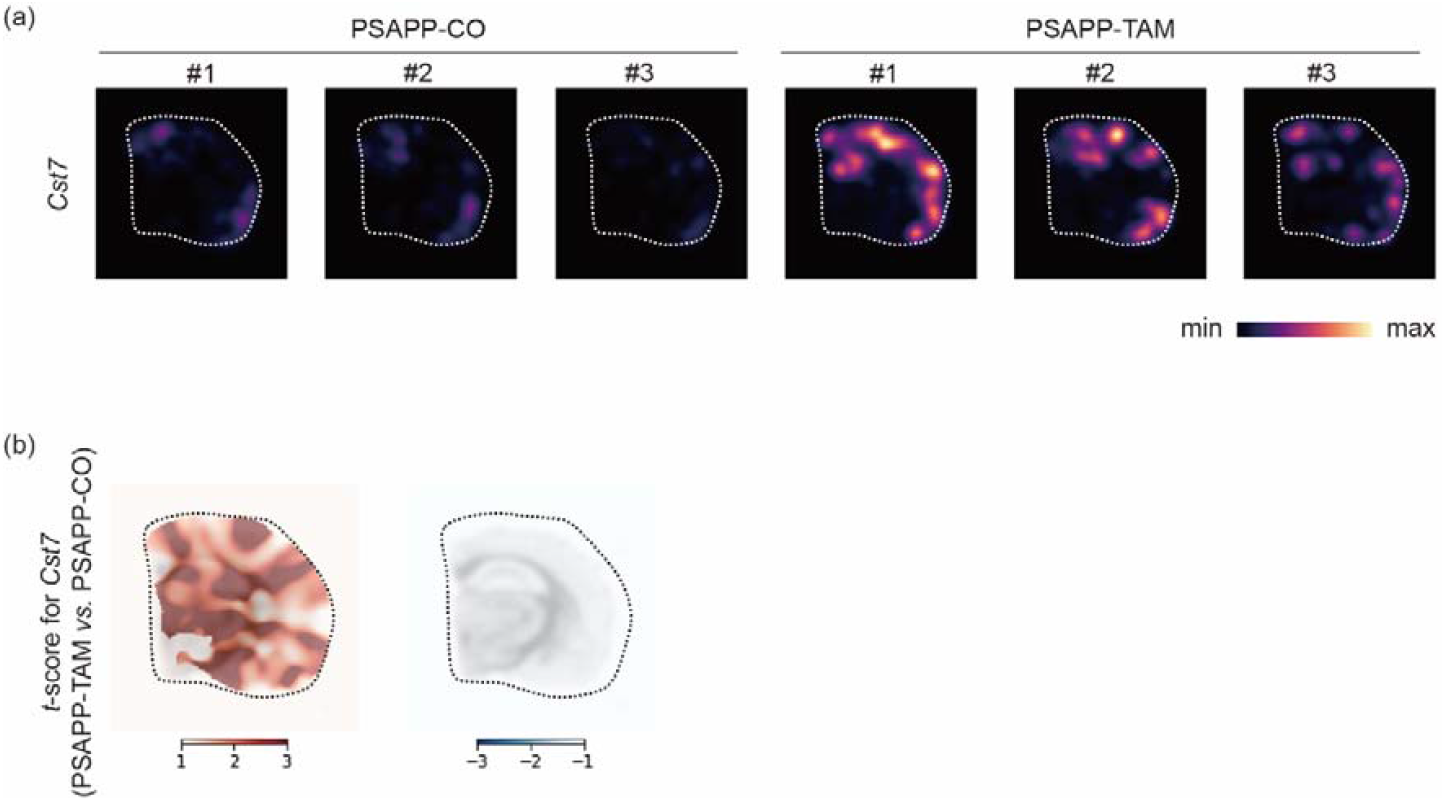
*Cst7* expression levels in spatially registered brain images of Alzheimer’s disease mouse model. (a) Heatmaps illustrated based on *Cst7* expression levels across two groups (PSAPP-CO and PSAPP-TAM). (b) Heatmaps illustrated based on the *T*-score estimated by pixel-wise T-test between two groups. **Abbreviation**: CO, corn oil; PSAPP, APP^KM670/671NL^/PSEN1^Δexon9^; TAM, tamoxifen;

In **Supplementary Fig. 2**, the representative parametric *T*-score maps were shown for the most significantly upregulated 30 genes in mice in PSAPP-TAM group, compared to those in PSAPP-CO group.

## Discussion

As ST technology becomes more ubiquitous, there is increasing demand to investigate the spatial gene expression patterns in complex organs such as the kidney, brain, intestines, and liver ^1,2^. Researchers acquire ST datasets from tissues exhibiting specific diseases or pathological conditions, detect alterations in gene expression within regions of anatomical interest and investigate their clinical implications. However, the lack of a methodology to align ST data anatomically for the various shape and coordinates of histologic tissues limits the direct comparison of gene expression patterns across various types of samples ^12^.

One of the main contributions of SpatialSPM is the development of a method that overcomes this challenge. Pixel-wise or voxel-wise statistical analysis, such as the well-established method called statistical parametric mapping ^13^, is commonly used for analyzing images with complex structures, such as the brain ^14^. SpatialSPM applies this method to transform ST data into image matrices, portraying every gene expression level of each ST dataset on a common template image, referred to as spatially registered 2D images. This process could facilitate direct comparison of gene expression patterns across similar regions in different histologic samples by pixel-wise statistical analyses, such as correlation analyses or T-tests, for specific target genes.

The **s**patial investigation of molecular change play a significant role in the analysis of disease pathophysiology and the mode of action of drugs ^8^. In this regard, SpatialSPM is a powerful technique that offers new insights into the underlying molecular mechanisms of various diseases, particularly in the brain. This is particularly relevant given the intricate relationship between anatomical structures and brain function. Furthermore, by utilizing cellular deconvolution for ST dataset ^15^, SpatialSPM empowers researchers to investigate the density of specific cell groups or pathway signatures. The cellular map can be inferred from the ST dataset and subsequently compared the groups using this proposed method.

However, some limitations to the use of SpatialSPM should be noted. Firstly, barcode-based ST technology provides sparse transcriptomic data with spots located apart from each other, rather than a complete pixel image or matrix. To address this issue, we generated 2D image matrices with continuous location information by assuming a Gaussian distribution of transcript data, followed by the spatial registration process. In this line, it is important to consider the technology platform used for generating ST dataset. It is necessary to expand the application and provide more detailed explanations of this method on various technology platforms, including image-based ST (such as Xenium, CosMX, and MERFISH) ^1^. Furthermore, it is important to consider a technical issue that arises during the tissue acquisition process for the ST dataset. For example, when analyzing brain tissue, the orientation or anatomical location of sectioning can result in completely different ST datasets. Such variations can introduce errors in the analysis process, potentially leading to false positive or false negative findings. Nevertheless, by employing well-structured histology and ST dataset, we believe that SpatialSPM can overcome these challenges and yield reliable results.

In conclusion, the development of SpatialSPM represents a significant advancement in the analysis of ST datasets, especially for enabling direct comparative analysis across tissue samples. This methodology leverages imaging analysis to study structurally important tissues, such as the brain. The capability to directly compare gene expression changes within the same specific anatomical region, even in the presence of variations in shape and spatial anatomy of histology, would facilitate a more comprehensive identification of disease-specific gene expression alterations. Furthermore, this capability would pave the way for the future development of targeted therapies.

## Materials and Methods

### Acquisition of ST dataset

Two ST datasets generated using the Visium platform technology (10x Genomics, CA, USA) were utilized for this study: the mouse intracranial hemorrhage model (GSE182127) ^9^ and the amyloid-depositing Alzheimer’s disease mouse model (*APP*^*KM670/671NL*^*/PSEN1*^Δ*exon9*^; PSAPP) with conditional *Inpp5d* knockdown in microglia (*PSAPP*/*Inpp5d*^fl/fl^/*Cx3xr1*^CreER+^) (GSE203424) ^11^. The gene expression matrix and metadata, including location coordinates, were loaded using scanpy version 1.6.1 ^16^. The expression numbers in the datasets were log-transformed after normalization by dividing the counts within each cell by the total number of transcripts, scaled by 10^6^.

### Generation of a two-dimensional image matrix of each gene expression based on the ST dataset

The multiple ST datasets, comprising each gene expression level data for every detection spot, were converted into 2D image matrices for every gene (**Fig. 1a**). We performed a convolutional operation, assuming that each gene expression value followed a Gaussian distribution based on its distance from the center of the spots. In detail, this calculation process imputed the expression level of the x-th gene at specific locations in the empty space between the spots by superimposing the Gaussian values of the gene expression levels based on the centers of the other neighboring spots (*i, j, k*, and so on) (**Fig. 1a**). The degree of the Gaussian distribution was specified to measure the count corresponding to a specific range (the value of the probability, 1-*alpha*) within a specific radius value. In this case, the radius was set to three times the distance between spots, and *alpha* was set to 0.01 for smoothing the gene expression patterns. This choice acts as a hyperparameter for Gaussian distribution-based gene expression values that were imputed between the spots. Afterward, the 2D image generated for each gene was compiled and loaded as a matrix to proceed to the next step.

### Spatial registration process of 2D image on a template image according to a transformation matrix

Initially, a score map of HVGs was constructed based on the generated 2D images from each ST dataset. This construction was achieved using the ‘*scanpy*.*tl*.*score_genes’* function in the Scanpy package (version 1.9.2) ^16^ by calculating every gene score (**Fig. 1b**). By matching the score map of HVGs, which was selected as the template image, with the other HVGs maps, a transformation matrix was generated. The transformation matrix was constituted by a non-linear transform with a combination of transformation methods. In detail, the implemented method includes four main steps: center-of-mass (COM) matching, translation, rigid, and affine transforms, and non-linear transform. Firstly, the COM matching step and the translation transform step align the images by estimating the translation parameters. The rigid transform step uses rotation and scaling to refine the alignment of the images. The affine transform step applies shearing and stretching to the images to further improve the registration. Finally, the non-linear transform step uses the symmetric diffeomorphic registration method to achieve a smooth deformation of the moving image. This step optimizes the spatial registration process by using the cross-correlation metric with Gaussian smoothing. The optimization was performed over multiple resolutions of the images. This transformation matrix was then applied to the matrices of all genes in the generated 2D images, resulting transformed matrices. This spatial registration process were performed using the ‘dipy’ package (version 1.4.1) ^17^ (**Fig. 1c**). Based on these transformed matrices, spatially registered 2D images were portrayed.

### Pixel-wise comparative analysis for the gene expression across the spatially registered 2D images

Comparative analysis at the pixel level was performed across the spatially registered images to evaluate gene expression levels in a specific anatomical ROI (**Fig. 1d**). This is because each specific pixel in the images was matched to the corresponding anatomical location. This study utilized ST datasets from two experimental animal brain models: the mouse intracranial hemorrhage model and the mouse model of Alzheimer’s disease, as working examples. In these two animal models, the levels of transcriptomes that changed in specific anatomical regions of the brain under specific conditions were compared to those of the control group, followed by statistical analysis.

In the mouse intracranial hemorrhage model, the statistical association between the concentration of treated Heme-Albumin and the transcriptome levels of genes was evaluated using Pearson correlation coefficients. In the mouse model of Alzheimer’s disease, pixel-wise independent T-tests were used to statistically compare the transcriptomes in the brains of PSAPP control mice with those of PSAPP mice with tamoxifen-sensitive microglial knockdown of *Inpp5d*.

### Data availability

The ST data in this study are available from the Gene Expression Omnibus (GEO) under the accession number GSE182127 and GSE20342.

## Acknowledgments

We thank all members of Portrai, Inc. for the discussion and technical support.

