## Supplementary Figures for "SpatialSPM: Statistical parametric mapping for the comparison of gene expression pattern images in multiple spatial transcriptomic datasets"

### Supplemental Figures

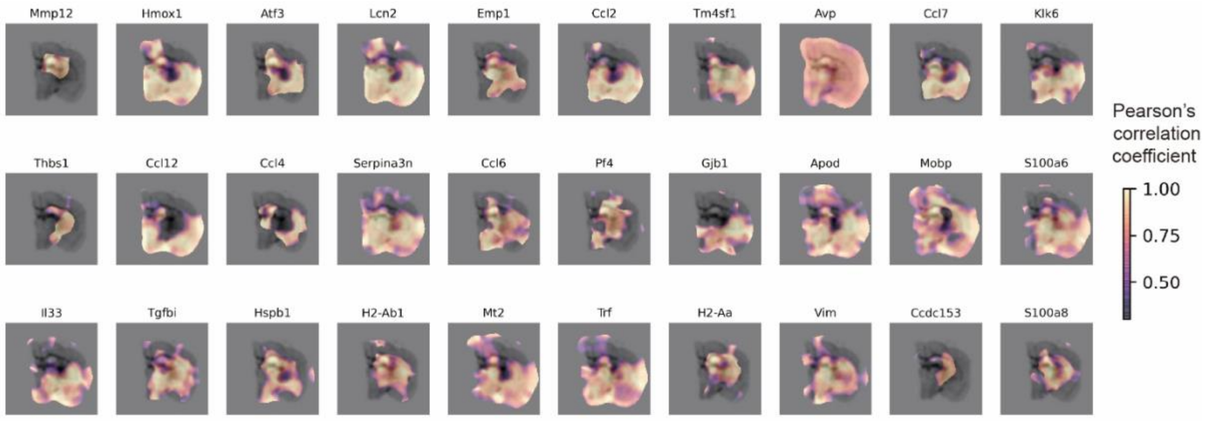


**Supplemental Figure 1. Representative parametric maps of genes highly correlated with Heme-Albumin concentration.** Top 30 genes highly correlated with Heme-Albumin concentration were presented in order (Pearson's correlation coefficient > 0.3).

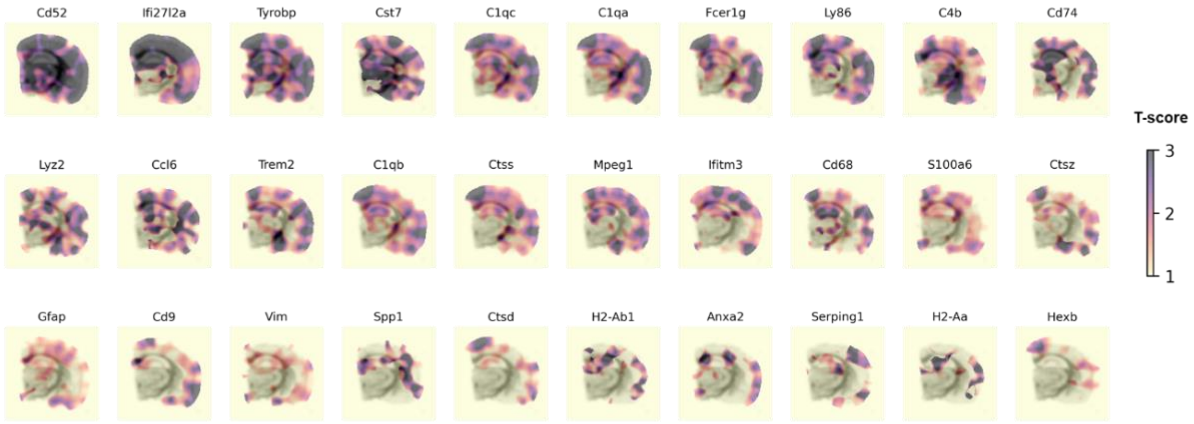


**Supplemental Figure 2. Representative parametric maps of genes upregulated in PSAPP-TAM mice.** The top 30 genes that were significantly upregulated in PSAPP-TAM mice compared to PSAPP-CO were presented (in order by *T*-scores).

**Abbreviation**: CO, corn oil; PSAPP, $APP^{KM670/671NL}/PSEN1^{\Delta exon9}$; TAM, tamoxifen;
